# Flight-induced compass representation in the monarch butterfly heading network

**DOI:** 10.1101/2021.04.07.438824

**Authors:** M. Jerome Beetz, Christian Kraus, Myriam Franzke, David Dreyer, Martin F. Strube-Bloss, Wolfgang Rössler, Eric J. Warrant, Christine Merlin, Basil el Jundi

## Abstract

For navigation, animals use a robust internal compass. Compass navigation is especially crucial for long-distance migrating animals like monarch butterflies, which use a sun compass to navigate every fall over 4,000 km to their overwintering sites. The central complex, a brain region equipped with sun-compass neurons, is proposed to control the butterfly’s heading. Although the activity of central-complex neurons exhibits a locomotor-dependent modulation in many insects, the consequences of such modulations for the coding of heading remain unexplored. Here, we developed tetrode recordings from tethered flying monarch butterflies to reveal how flight modulates heading representation. We found that during flight, heading-direction neurons change their tuning, transforming the central-complex network to function as a global compass. This compass coding is characterized by the dominance of steering feedback and allows for robust heading representation even under unreliable visual scenarios, an ideal strategy for maintaining a migratory heading over enormous distances.

## Introduction

Goal-directed navigation requires animals to constantly register their heading in space. This is of particular importance during long-distance migration, as exhibited by many vertebrates, such as birds and bats ^1^. Similarly, monarch butterflies (*Danaus plexippus*) annually migrate from southern Canada and the northern United States to their overwintering habitat in central Mexico ^2,3^. To maintain proper migratory direction, monarch butterflies rely on an internal compass that integrates multimodal orientation signals, including the sun ^4,5^. Sun compass information is processed in a brain region termed the central complex ^6,7^. In this brain region, compass neurons are thought to encode the butterflies’ heading, like head-direction cells in vertebrates ^8–12^. Data from the central complex of *Drosophila* show that heading is topographically represented in a neural network composed of E-PG cells ^13–15^. Within the E-PG network, heading is represented by a single activity bump that changes its position when the insect changes its heading direction ^13,16^. The activity bump in E-PG cells results from the integration of visual signals and mechanosensory information ^14,17–19^, as well as idiothetic, self-motion signals ^13,20,21^. Although the multimodal input may explain locomotor-dependent modulations in the activity of central-complex neurons ^22,23^, the function of these state-dependent modulations is still poorly understood. To shed light on this, we investigated how the locomotory state affects sun compass coding in the monarch butterfly.

In the past, due to methodological limitations, butterfly’ sun compass neurons have only been recorded in restrained animals ^6^. Here, using monarch butterflies, we investigated how the heading-direction network operates when the animal is free to steer with respect to a sun stimulus. We observed that when the animals were free to steer during flight, the heading-direction network modified its angular tuning, i.e. remapped from a vision-based sun-bearing coding found in quiescent animals to a compass coding during flight. The transformation to a compass was reflected by a state-dependent change in angular sensitivity and tuning. At flight onset, a substantial number of neurons started to encode the solar azimuth, while others ceased to encode the sun position, a process that depends on changes in octopamine levels. Changes in angular tuning were accompanied by a state-dependent switch in cue hierarchy between visual and idiothetic inputs. Importantly, the heading-direction network transformed into a compass that was dominated by idiothetic cues when the tethered flying butterflies were allowed to perform actual turns. The dominance of idiothetic cues allowed a robust heading representation even under ambiguous or conflicting visual settings. Thus, in the monarch butterfly central complex, flight activity shapes the heading-direction network into a multimodal, global compass that allows these insects to reliably maintain their migratory southward direction.

## Results

### State-dependent changes in angular sensitivity of central-complex neurons

To perform tetrode recordings from tethered flying butterflies, we designed an indoor flight arena (Fig. 1A). The arena’s upper inner circumference was equipped with a strip of individually switchable green light spots serving as sun stimuli that butterflies could use for orientation. During neural recordings from the central complex of tethered butterflies centered in the arena (Fig. 1B, Supplementary Figure 1), an optical encoder measured the animal’s heading direction. First, we recorded from quiescent butterflies that rested on a platform at a fixed heading (0°; *quiescent pre-flight*; Supplementary Movie 1) and that were stimulated by moving the sun stimulus along a 360° circular path around the animal. To investigate the influence of proprioceptive effects from flight activity on the neural response, we then presented the same sun stimulus, but now with the butterfly flying at a fixed heading (0°; *fixed flight*; Supplementary Movie 2). A subset of central-complex neurons (66 of 150 recorded neurons; 44%) exhibited state-dependent angular tuning to the sun stimulus. During *fixed flight*, 50 neurons (33%; 8 + 29 + 13 neurons outside the gray area in Fig. 1C) encoded the solar azimuth as indicated by an increase in firing rate at neuron-specific azimuthal positions of the sun stimulus (Rayleigh test: p < 0.05). In contrast, during quiescence, the same neurons showed stable firing rates across different sun positions (Rayleigh test: p > 0.05) and therefore did not encode the solar azimuth (Fig. 1D, green dataset). Conversely, 16 neurons (11%, 2+14 neurons of gray area in Fig. 1C) showed angular sensitivity during quiescence but ceased to encode the solar azimuth during *fixed flight* (Fig. 1D, blue dataset). The remaining 84 of the 150 recorded neurons (56%) did not change their angular sensitivity to the sun stimulus during fixed flight: 76 neurons were sensitive (51%; 56 + 20 neurons in merged gray and blue areas of Fig. 1C) and 8 neurons insensitive (5%; green area in Fig. 1C) during both locomotory states. To test whether changes in angular sensitivity reflect recording instability, e.g. subtle electrode movements induced by flight, we allowed the butterfly to settle on the platform after flight and retested the neural sensitivity in 63 neurons (*quiescent post-flight*). All 63 neurons retained their *quiescent pre-flight* angular sensitivity (example shown in Fig. 1D), suggesting that changes in angular sensitivity observed during flight depended solely on the animal’s locomotory state. We also excluded that we recorded from different neurons across the behavioral states by assessing the spike-shape stability. This was done by computing the averaged spike shapes from each neuron and locomotory state (i.e., *quiescent pre-flight*, *fixed flight*, and *quiescent post-flight*). These averaged spike shapes were correlated with each other, and the correlation coefficients were statistically compared with the ones obtained by correlating neuron-specific spike shapes with averaged spike shapes of randomly chosen neurons (Fig. 1E–1F, cf. Online Methods). For all neurons of the present study, the averaged spike shapes within spike-sorted neurons statistically correlated better than with averaged spike shapes across different neurons. This strongly suggests that we recorded from the same neurons throughout the entire experiment.

**Fig. 1.**
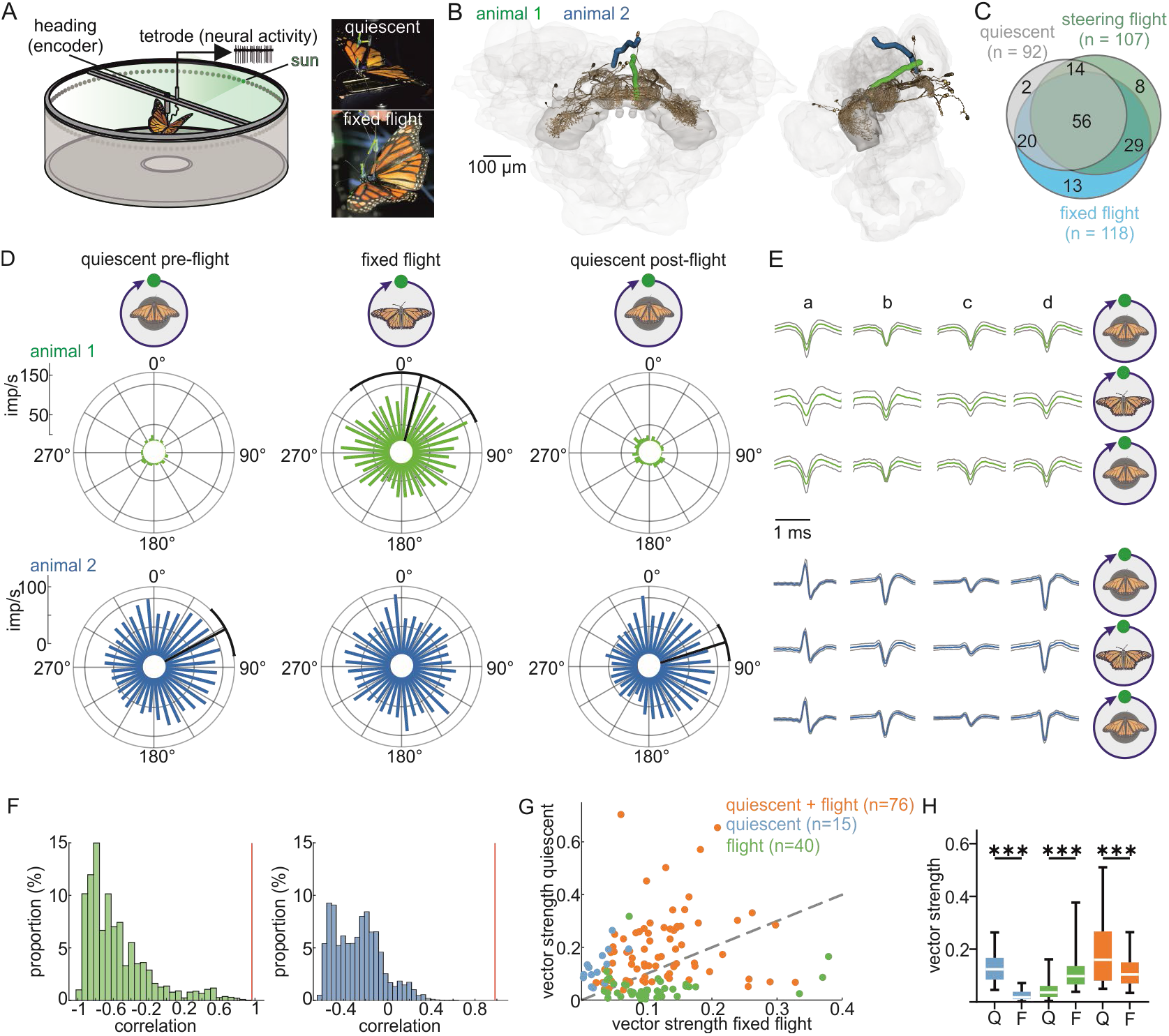
State-dependent change in angular sensitivity in butterfly central-complex neurons. **A**, Schematic drawing of the setup. **B**, Anterodorsal (*left*) and lateral (*right*) view of reconstructed tetrode tracks from two animals (*blue and green*) warped into the standardized monarch butterfly central complex ^7^ including prominent central-complex neurons ^7^ (*brown*). **C**, Number of neurons showing angular sensitivity to the sun stimulus across different behavioral states, i.e. *quiescent, fixed flight, steering flight*. **D**, Responses of two example neurons (*blue*, *green*, same color code as in B) to the moving sun stimulus (animals faced 0°) in different behavioral states, i.e. *quiescent pre-flight (left), fixed flight (middle), quiescent post-flight (right)*. *Black line* and *arc* represent the preferred firing direction ± circular SD of neurons that encoded the solar azimuth (Rayleigh test: p < 0.05). **E**, Spike shapes (*mean ± IQR*) of individual tetrode channels (*a*-*d*) for each behavioral state for the *blue* and *green* neurons shown in (D). **F,** Comparison of the spike shape correlation (for the *blue* and *green* neuron in D) between two behavioral states (*red vertical line*) and a modelled distribution of spike correlations. **G,** Comparison of vector strengths calculated for *quiescent* and *fixed flight*. Neurons that showed angular sensitivity exclusively during *fixed flight* and *quiescence* are shown in *green* and *blue*, respectively. Neurons encoding the solar azimuth in all locomotion states are indicated in *orange*. **H**, Statistical comparison of vector strengths across locomotion states (Q = *quiescent*; F = *fixed flight*; Wilcoxon signed rank test: *** = p < 0.001).

Next, we examined how state-dependent changes in angular sensitivity are reflected in the broadness of neural tuning. For each neuron and locomotory state, we calculated the neuron’s vector strength. The higher the vector strength, the narrower the neural tuning, and vice versa. As expected, neurons that started to encode the solar azimuth during fixed flight (Rayleigh-test: p < 0.05) exhibited higher vector strengths during *fixed flight* compared to *quiescence* (Wilcoxon signed rank test: p < 0.0001; green data points in Fig. 1G–1H). In contrast, neurons that ceased to encode the solar azimuth during *fixed flight* (Rayleigh test: p > 0.05) had lower vector strengths during *fixed flight* compared to *quiescence* (Wilcoxon signed rank test: p < 0.0001; blue data points in Fig. 1G–1H). Neurons encoding the solar azimuth irrespective of the animal’s locomotory state showed broader tuning during *fixed flight* compared to *quiescence* (Wilcoxon signed rank test: p < 0.0001; orange data points in Fig. 1G–H).

### Remapping of the heading-direction network during steering flight

To test if changes in heading coding occurred when tethered flying butterflies were free to rotate around their yaw axis while the sun stimulus was set at 0° (*steering flight*; Supplementary Movie 3). We classified 70 of the 150 recorded neurons as encoding the solar azimuth during *quiescence* and *steering flight* (Fig. 2A–C; 47% of the neurons; 14 + 56 neurons of gray portion in Fig. 1C). While some neurons showed a similar angular tuning across *quiescence*, *fixed*, and *steering flight* (magenta neuron in Fig. 2A–C), many others shifted their preferred firing direction, during *steering flight* compared to *quiescence* (Fig. 2C, black lines in circular plots of blue neuron; Supplementary Figure 2), suggesting a remapping of the neural network. When comparing the neural response across a population of 27 neurons that were sensitive at each behavioral state, we found that the remapping mainly occurred during *steering flight* (Fig. 2D) but rarely between *quiescence* and *fixed flight*, and *quiescence pre-* and *post-flight* (Fig. 2E and 2G; respectively, n = 76 and 46 for *quiescent* versus *fixed flight* and *quiescent pre-* versus *quiescent post-flight*; V-test: p < 0.0001). The shifts in angular tuning during *steering flight* were arbitrary across neurons, indicated by a uniform distribution (Fig. 2F; Rayleigh test: p = 0.09, n = 70). While, during flight, the relative angular position between butterfly and sun stimulus dynamically changed with the butterfly’ heading direction, stimulus history and velocity were constant in quiescence. To exclude the possibility that the observed remapping resulted from differences in the spatiotemporal dynamics of the stimulus, we measured the neural tuning when the sun stimulus unpredictably moved at different directions and velocities in quiescence (Supplementary Movies 4 and 5; Supplementary Figure 3A-C; n = 42 neurons). We found that the neural tuning was neither affected by stimulus history nor stimulus velocity (V-tests: p < 0.001). Remapping could also be evoked by the coding of turns ^20,21,24^, which may obscure angular tuning to our sun stimulus. Only 7 (5%) of our 150 recorded neurons conjunctively encoded flight turns and heading (example neuron in Supplementary Figure 3D-E), confirming that coding of angular turns was not responsible for remapping. Taken together, our data suggest that the remapping in the heading-direction network mainly occurs during *steering flight* and is dependent on the animal’s locomotory state.

**Fig. 2.**
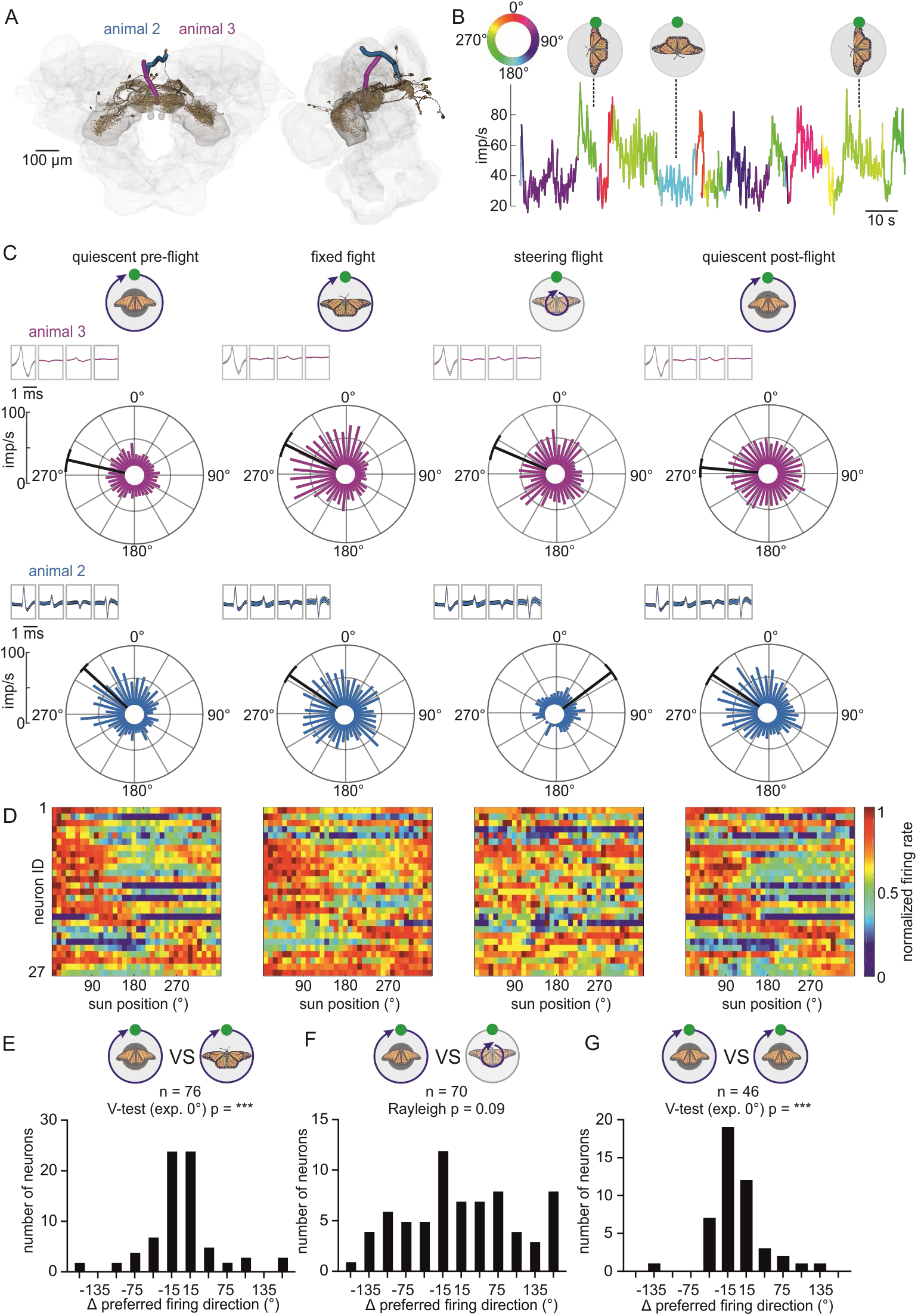
State-dependent changes in angular tuning of butterfly central-complex neurons. **A**, Reconstructed tetrode tracks one from each animal, indicated in different colors. **B**, Neural firing rate from a central-complex neuron as a function of the animal’s heading coded in color. The neuron increased its firing rate when the animal was heading at 270° while neural activity decreased when the animal was heading at 180°. **C,** Firing rates of two neurons (*magenta*, *blue*) at different sun positions relative to the animal’s heading (pointing towards 0°) for four behavioral states. Corresponding spike shapes (*mean ± IQR*) are indicated at the top left corner of each circular plot. **D,** Neural activity (color coded) of 27 neurons (*y-axis*) plotted against the sun position (*x-axis*) for each behavioral state. Only neurons that were sensitive (Rayleigh test: p < 0.05) across all four behavioral states are included. For all neurons that were sensitive during *quiescence* and *steering flight* see Fig. S3. **E-G**, Changes in preferred firing direction between *quiescent pre-flight* and *fixed flight* (E), *quiescent pre-flight* and *steering flight* (F), and *quiescent pre-flight* and *quiescent post-flight* (G).

### A 360° compass of heading angles is computed during flight

While in *Drosophila*, remapping of the heading-direction network could be induced by presenting different visual scenes ^14,15^, a state-dependent remapping was not found ^14^. In contrast to this, we here demonstrate for the first time a state-dependent remapping in the heading-direction network of a long-distance migrating insect. To understand the biological role of state-dependent remapping in monarch butterflies, we compared the distribution of preferred firing directions during *quiescence* with the distribution during *steering flight*. During *quiescence*, the neurons predominantly encoded the butterfly’s frontal hemisphere (n_quiescent_ = 92 neurons; Fig. 3A; V-test: p < 0.001), while during *steering flight* the neurons fully tiled 360° representation of heading (n = 107; Fig. 3B; Rayleigh test: p = 0.83). These results strongly suggested that the heading-direction network changes its coding strategy and only functions as a compass when the butterfly is flying.

**Fig. 3.**
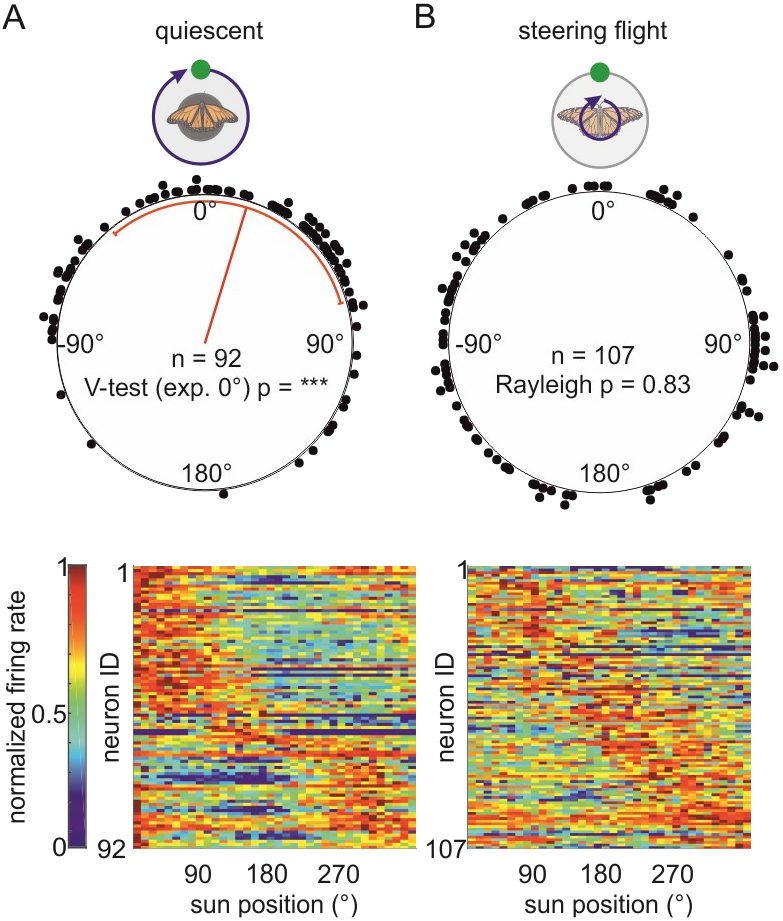
State-dependent transition of the butterfly central complex to a compass. **upper A-B,** Distribution of the preferred firing directions during *quiescence* (A) and *steering flight* (B). The preferred firing directions were biased towards the frontal visual field during *quiescence* while they showed a uniform 360° distribution during *steering flight*. **lower A-B**, Neural activity (color coded) of respectively 92 neurons and 107 neurons plotted against the sun position (*x-axis*) for quiescence (*left heatmap*) and steering flight (*right heatmap*). Differences in the number of neurons in each colormap come from the fact that only neurons that were sensitive (Rayleigh test: p < 0.05) to the respective locomotion-state were considered.

To characterize the coding strategy of the heading-direction network across different locomotory states flight, we performed experiments where we tested the neural coding of the same neurons in quiescent and flying butterflies but from butterflies presented with an ambiguous visual stimulus i.e., two suns set 180° apart (2-sun experiment). When the two suns were rotated by 360° around a quiescent butterfly (Fig. 4A; *quiescent pre-flight* in Fig. 4B–C), heading-direction neurons showed a bimodal response reflecting the 180° periodicity of the visual stimulus. This bimodal response was consistent across each full stimulus rotation (middle column in Fig. 4B) indicating that the heading-direction network of quiescent butterflies equally weighted and computed both suns as isolated visual cues. In contrast, the same neurons represent the visual scene by a unimodal response during steering flight (Fig. 4A; *steering flight* in Fig. 4B–C). Thus, the network forms a compass that globally integrates both suns into a 360° visual scene and allows the butterfly to encode an explicit heading. Importantly, as soon as the animal settled (*quiescent post-flight*), the heading-direction network only receives visual information and therefore exhibited a 180° periodicity in heading coding, similar to the coding observed prior to flight (Fig. 4B–C). State-dependent transitions between unimodal and bimodal responses were highly significant (Fig. 4D; RM one-way ANOVA + Holm Sidák’s multiple comparison test: p < 0.001, n = 33 neurons). Together, these results demonstrate a state-dependent functional remapping in the central complex: In quiescence, the heading-direction neurons represent heading in a visual-based sun bearing coding. In contrast, the same neurons transform into a compass coding during flight, which allows butterflies to maintain all possible heading directions in a body invariant manner.

**Fig. 4.**
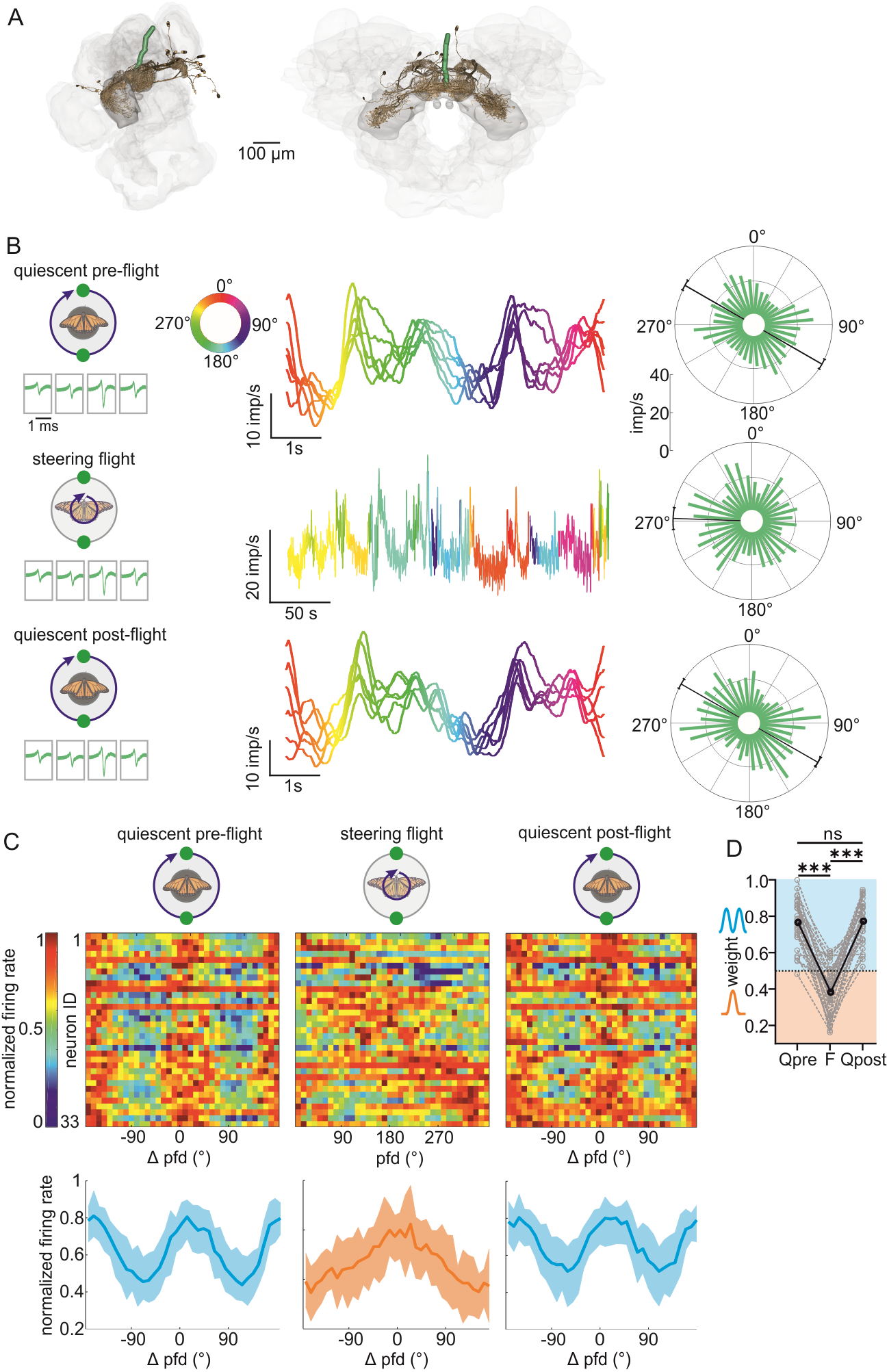
Heading coding of central-complex neurons under an ambiguous visual scene. **A**, Tetrode position of the neuron recorded in (B). **B**, The neural response to the 2-sun stimulus in quiescent (stimulus was rotated 6 times around the animal; *first and third row*) and flying monarch butterflies (static stimulus; *second row*). (*Left*) Average spike shapes are shown below the schematic icons. (*Middle column*) Neural response to individual stimulus rotations (sun azimuth is color coded) in *quiescence* and the response of the same neuron during flight (animal’s heading color coded). (*Right*) Circular plots summarize the neural tuning for the respective locomotion state. **C**, (*Top*) Responses of 33 neurons in each behavioral state. Along the x-axis, the heatmaps are centrally aligned to the neuron’s preferred firing direction (*pfd*) in the *quiescent pre-flight* condition. Along the y-axis, the neurons are ordered according to the pfd measured during steering flight. (*Bottom*) Averaged response (mean ± quantile) of all 33 neurons during quiescence (*left*, *right*) and during flight (when the neurons’ maximum responses were shifted to 0°). **D**, Weights of the responses of the 33 neurons to a bimodal or unimodal model for each behavioral state, i.e. *quiescent pre-flight (Q_pre_), steering flight (F), quiescent post-flight (Q_post_)*. A weight between 0 and 0.5 indicates that the neural response was unimodal, whereas a weight between 0.5 and 1 indicates that the neural response was bimodal. Mean weights are indicated by black lines. Friedman test: *** = p < 0.0001; ns = p > 0.05.

### Octopamine changes angular sensitivity

What mechanisms underly the state-dependent transformation into a sun compass during flight? As some neurons increased their firing rates during flight (e.g., Fig. 1D, green neuron), we wondered whether changes in neural firing rate contribute to the state-dependent changes in angular sensitivity. We found that this was not the case as changes in neural firing rates during quiescence (pre- & post-flight) and flight (Fig. 5A) were not sufficient to induce any state-dependence changes (orange data points in Fig. 5A). In flies, the neuromodulator octopamine, the invertebrate analog of noradrenaline ^25^, is expressed during flight and has been linked to state-dependent changes in the neural sensitivity of optic lobe and descending neurons ^26,27^. To test if octopamine could mediate the state-dependent changes we observed in the heading-neural network, we topically applied chlordimeform (CDM), an octopamine agonist, onto the brain of quiescent butterflies (N = 13 animals; n = 50 neurons) to mimic potential flight-induced physiological effects ^26,28^. Following CDM application, eight neurons (15 %, Fig. 5B–C, cyan neuron; neuron ID 17-24 in Fig. 5D) started to encode the solar azimuth, whereas 16 neurons (31 %, Fig. 5B–C, magenta neuron: neuron ID 1-16 in Fig. 5D) lost angular sensitivity. For the remaining 26 neurons (51 %), we could not detect an effect of CDM on angular sensitivity. The vector strength of neurons from CDM-treated *quiescent* butterflies was similar to that for neurons of non-treated butterflies during *fixed flight* (compare magenta /cyan data points in Fig. 5E–F and blue/green data points in Fig. 1G–1H). While flight-induced changes in angular sensitivity could be mimicked by CDM treatment, CDM did not however induce any remapping (Fig. 5G, V-test: p < 0.001). These results suggest that idiothetic cues exclusively processed during steering flight are responsible for remapping the butterflies’ heading-direction network into a sun compass.

**Fig. 5.**
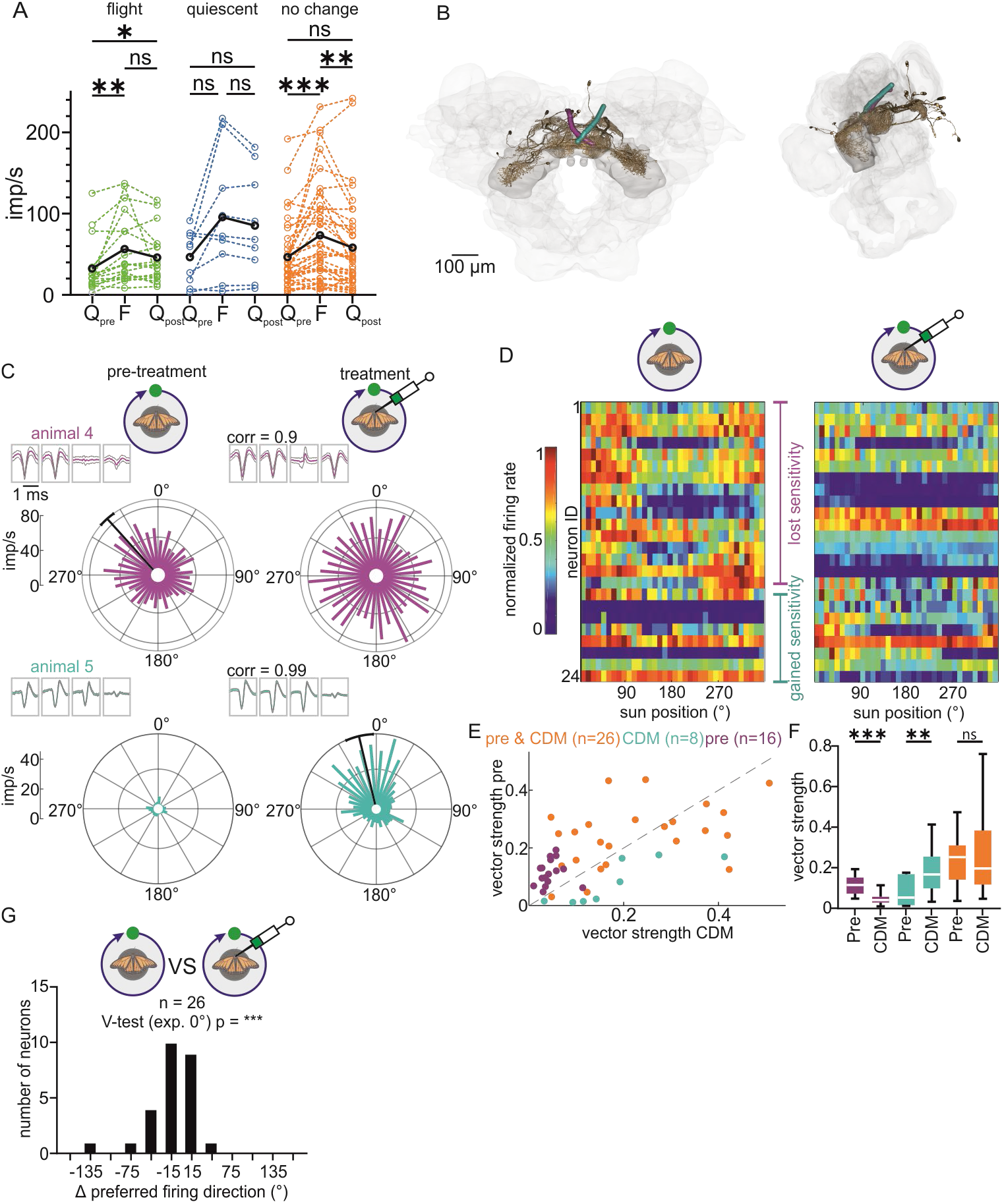
Changes in angular sensitivity after chlordimeform (CDM) application. **A**, Mean firing rates at different locomotion states, i.e. *quiescence* (pre: Q_pre_ and post: Q_post_) and *fixed flight* (F) for three different neuron subpopulations. *Green neurons* (n = 50) were sensitive only during *fixed flights* while *blue neurons* (n = 16) were sensitive only during *quiescence*. *Orange neurons* (n = 76) were sensitive in both locomotion states. *Black lines* represent mean values for each subpopulation. Friedman test: *** = p < 0.001; ** = p < 0.005; * = p < 0.05; ns = p > 0.05. **B**, Tetrode tracks of the neurons shown in C. **C**, Circular plots of two neurons recorded from two butterflies [color code as in (B)] that changed their sensitivity after CDM application in quiescent animals. Corresponding spike shapes (*mean ± IQR*) and correlation coefficients (*corr*) are shown at the top left corner of each circular plot. **D**, Activity of 24 neurons (*Left*: tuning before application; *right*: tuning after application) that gained/lost angular sensitivity to the sun stimulus after CDM application. **E,** Comparison of vector strengths before (*pre*) and after CDM application. *Cyan* and *magenta* dots represent neurons that showed angular sensitivity exclusively after or before application. Neurons encoding the solar azimuth irrespective of CDM application are shown in *orange*. **F,** Statistical comparison of vector strengths before (*pre*) and after CDM application (Wilcoxon signed rank test: *** = p < 0.001; ** = p < 0.005). **G**, Preferred firing directions were stable after CDM application.

### Integration of idiothetic and visual inputs induces remapping during flight

During steering flight, insects receive a multitude of different idiothetic signals, including proprioceptive signals from flight muscles ^29^, vestibular-like signals ^30^, and motor efference copies. Vestibular-like signals inform animals about rotations and provide sufficient information to encode the insect’s heading even when the animals orient in darkness ^13,30^. To test the relevance of vestibular-like signals for heading coding in monarch butterflies, we mounted the butterflies on a rotation stage in the arena’s center. This allowed us to rotate the butterflies by 360° in the presence of a static sun stimulus (*quiescent rotation*; Fig. 6A and Supplementary Movie 6) followed by a rotation in darkness (*darkness rotation*). Similar to results in cockroaches ^30^ and rodents ^31,32^, in these conditions, the preferred firing directions of the butterfly neurons were invariant to the presence of visual stimuli (Fig. 6B–E; n = 16 neurons; V-test: p < 0.0001). The persistence of the neural tuning in darkness shows that the neurons represented heading directions in quiescence. The neural tuning during *darkness rotation* was broader than during *quiescent rotation* (Wilcoxon signed rank test: p = 0.0076; Fig. 6D and Supplementary Figure 4), consistent with the fact that computing heading directions only with idiothetic cues accumulates errors over time ^10^. The preferred firing directions measured in the presence (*quiescent rotation*) and absence (*quiescent*) of vestibular-like signals were almost identical (Fig. 6D and 6F; n = 16 neurons; V-test: p < 0.0001). Therefore, vestibular-like signals alone did not account for remapping of the heading-direction network as observed in the same neurons during *steering flight* (Fig. 6D and 6G; n = 16 neurons; Rayleigh test: p = 0.6). Alternatively, steering commands in the form of motor efference copies and reafference signals might trigger remapping. Each motor command is internally stored and compared with reafferent signals, representing sensory feedback associated with turns ^33–36^. This closed loop, from here on referred to as an “efference-reafferent loop”, allows the animal to predict sensory changes associated with turns. To test the influence of the “efference-reafferent loop” on heading coding, we tethered the butterflies and suspended them to a rotation stage that allowed us to subject them to 360° rotations and thus control their heading during fixed flights (Fig. 6H; *rotation fixed flight*; Supplementary Movie 7). We found that even in the presence of vestibular-like and proprioceptive signals in the absence of a functional “efference-reafferent loop”, the heading-direction network of butterflies performing fixed flights showed the same angular tuning as during *quiescence* (Fig. 6I–6J; n = 17 neurons; V-test: p< 0.001). Thus, our results indicate that a proper “efference-reafferent loop” is essential to transform the monarch butterfly heading-direction network into a compass during steering flight.

**Fig. 6.**
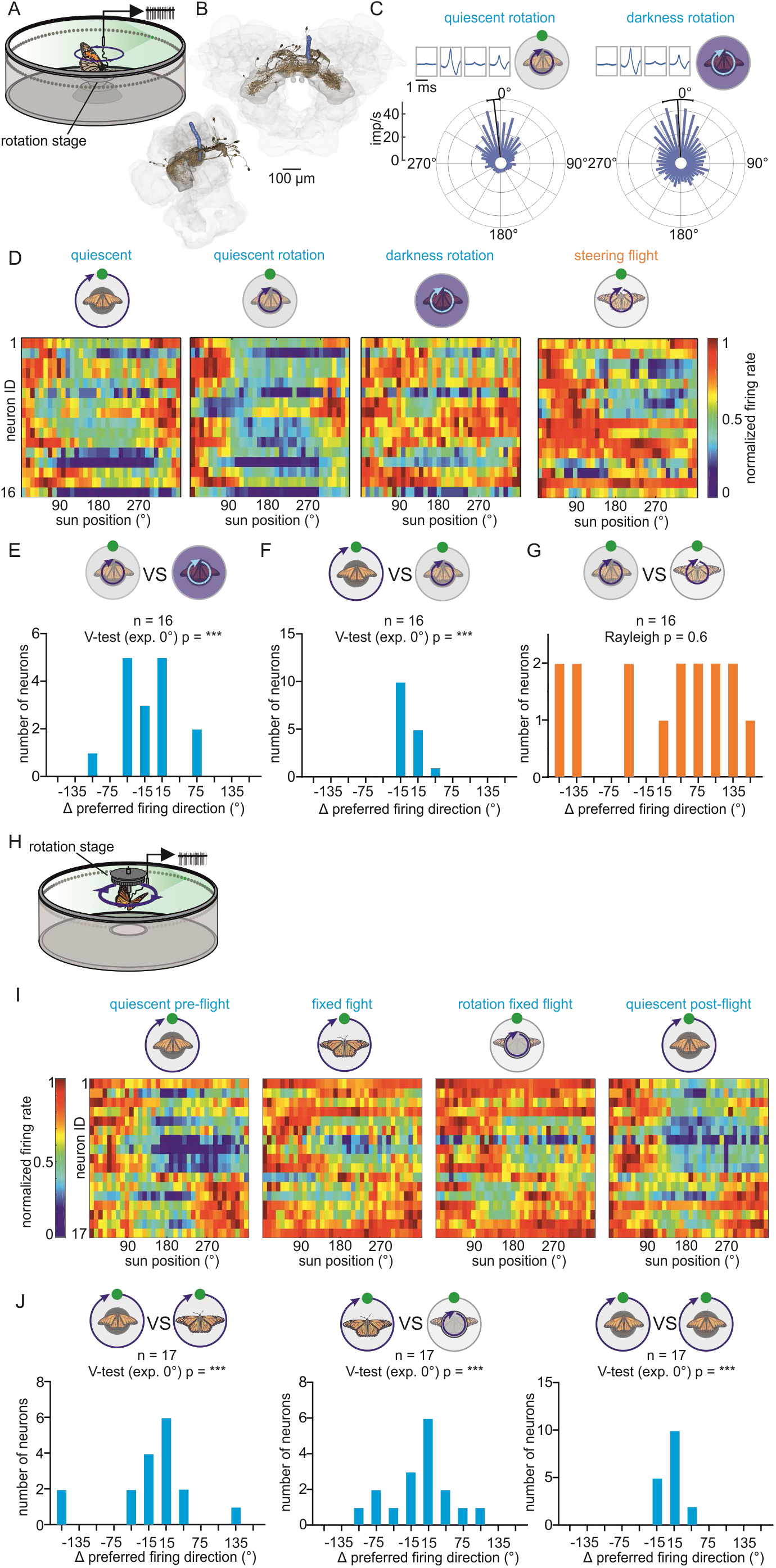
Motor efference copies and reafference signals induce remapping of the heading-direction network during flight. **A**, Illustration of the setup to test for the influence of vestibular-like signals. Angular sensitivity was measured while rotating a quiescent butterfly on a rotation stage. **B**, Tetrode position during the experiment in which the example neuron in (C) was recorded. **C**, A neuron showing a preferred firing direction when the butterfly was rotated in the presence of the sun stimulus (*left*) and in darkness (*right*). **D**, Neural activity from 16 neurons in each behavioral state is plotted against the sun position. Only neurons that were sensitive (Rayleigh test: p < 0.05) in all behavioral states are included. **E-G**, Changes in the preferred firing direction during *quiescent rotation* and rotation in darkness (*darkness rotation*) (E); in the absence (*quiescent*) and in the presence of vestibular-like input (*quiescent rotation*) (F); and *quiescent rotation* and *steering flight* (G). **H**, Illustration of the setup to test for the influence of vestibular-like and proprioceptive signals. Angular sensitivity was measured while rotating a suspended tethered butterfly that was allowed to perform flight but not to steer (*rotation fixed flight*). **I**, Neural activity of 17 neurons for each behavioral state, i.e. quiescent pre-flight, fixed flight, rotation fixed flight, and quiescent post-flight is plotted against the sun position. **J**, Changes in preferred fight direction between quiescent pre-flight and fixed flight (*left*), quiescent pre-flight and rotation fixed flight (*middle*), and quiescent pre-flight and quiescent post-flight (*right*).

### Idiothetic cues processed during steering flight dominate over visual cues

The explicit heading representation observed during flight in our “2-sun experiment” (Fig. 4) might be established by a hierarchy of cues during steering flight. To examine which sensory input, i.e. visual or idiothetic, is the most relevant for the heading-direction network in different behavioral states, we performed experiments in which both inputs were set in conflict in quiescent and steering flying butterflies. This was accomplished by displacing the sun stimulus by 180° during the experiment. When displacement occurred in quiescent rotating butterflies, the neural tuning shifted with the sun by about 180° (Fig. 7A–B; n = 37 neurons; V-test: p < 0.001). These results show that the coding of heading is dominated by the sun stimulus when quiescent animals are rotated when these receive vestibular-like inputs (see Fig. 6C–E). Interestingly, when we repeated the same experiments with steering flying butterflies, the neural tuning did not shift after stimulus displacement (Fig. 7C–D; n = 22 neurons; V-test: p = 0.02 with 0° as expected value). Thus, although the butterflies had the same stimulus setting during *quiescence* and *steering flight*, we demonstrate the existence of a state-dependent switch in the cue hierarchy in monarch butterflies. While the sun position dominated the heading coding in quiescence, idiothetic cues such as the “efference-reafferent loop” were ranked higher than the sun during flight.

**Fig. 7.**
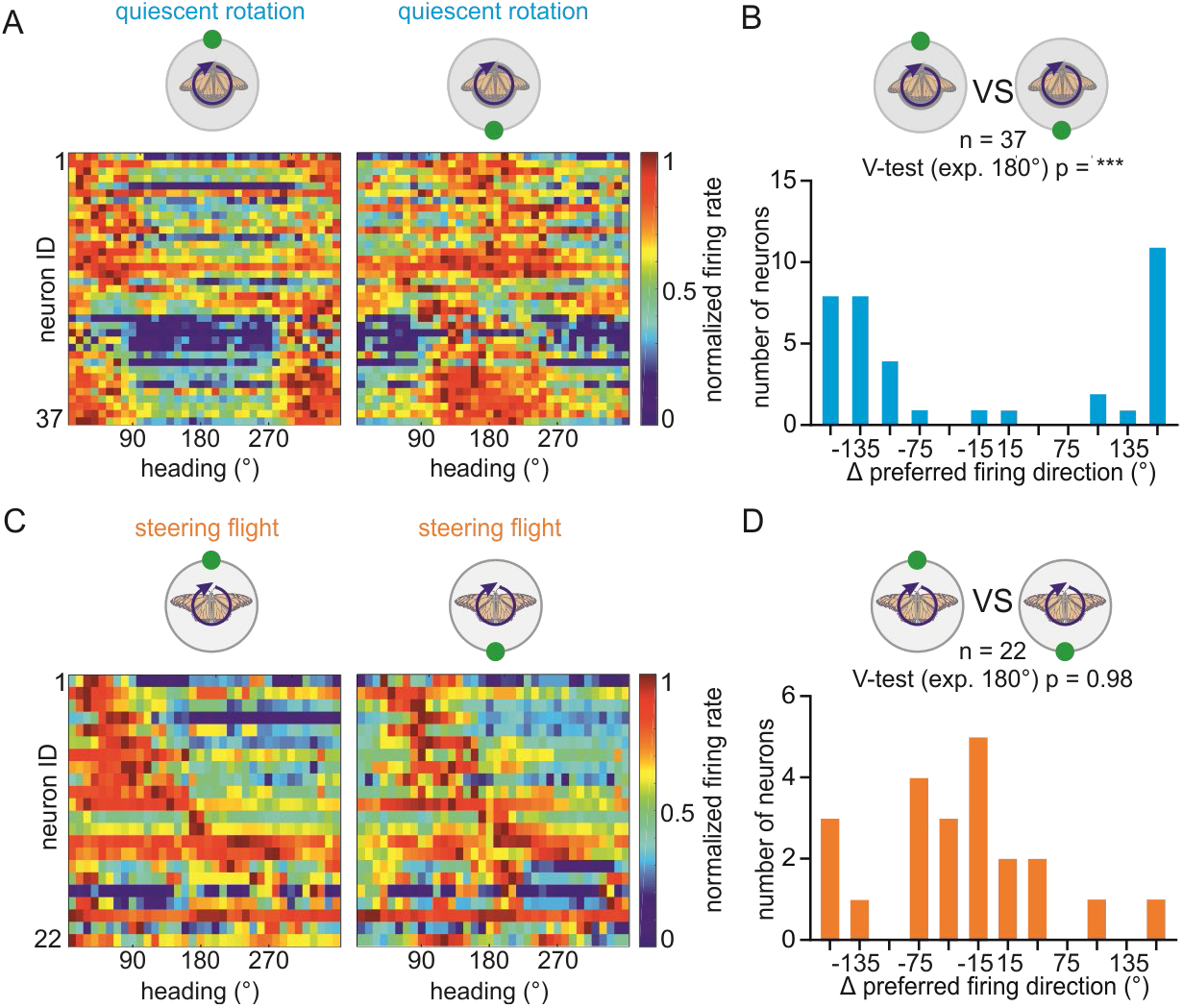
Heading coding is dominated by visual cues during quiescence while idiothetic cues dominate during flight. **A,** The neural activity of 37 neurons plotted against the animal’s heading when the sun stimulus was positioned at 0° (*left*) and 180° (*right*) and the animal was rotated by 360° around its body axis (*rotation*). **B,** Changes in the preferred firing direction between the sun stimulus positioned at 0° and at 180° when the animal was rotated. Most neurons shifted their preferred firing directions by 180° (V-test: p < 0.001). **C,** The neural activity of 22 neurons plotted against the animal’s heading when the sun stimulus was positioned at 0° (*left*) or at 180° (*right*) while the animals were able to steer during flight. **D,** The changes in preferred firing direction between the sun stimulus positioned at 0° and at 180°. The change of preferred firing direction was clustered around 0°.

## Discussion

In the present study, we show substantial state-dependent effects in the heading-direction network of the monarch butterflies, including flight-induced octopaminergic changes in angular sensitivity and steering-induced changes in angular tuning. We provide the first evidence in insects that actual steering has a strong impact on the processing of heading information in the brain, as previously shown in mammals ^31,37^, and demonstrates the biological relevance of state-dependent changes in the central complex. Depending on the locomotory state, the heading-direction network switches from a sun-bearing coding in *quiescence* to a proper sun compass coding during flight. A compass tiling 360° of azimuth allows a robust representation of the butterfly’ heading even under ambiguous (2-sun experiment) or variable (conflict experiment) visual settings. Robust heading representation is essential for migratory butterflies as they must maintain a stable flight course over an enormous distance while travelling through diverse visual scenes.

### Changes in angular sensitivity of heading-direction neurons

While recent studies reported state-dependent sensitivity changes of central-complex neurons ^22,23,38^, the neural mechanisms underlying these changes have remained elusive. State-dependent changes in more peripheral brain regions like the optic lobes of blowflies ^26^ and *Drosophila* ^39–41^ have been attributed to a flight-induced increase in the octopamine levels. Our results suggest that octopamine contribute to flight-induced changes in angular sensitivity of central-complex neurons. Unlike what has been shown in visual neurons of fruit flies ^41^, changes in angular sensitivity did not correlate with a state-dependent change in the firing rate. In *Drosophila*, it has been reported that ring neurons sending visual information into the E-PG network alter their sensitivity to visual cues in a state-dependent manner ^14,17^, making this cell type a possible neuronal substrate for the state-dependent changes in angular sensitivity found in monarch butterflies. By modulating the visual input from the population of ring neurons in a state-dependent manner, the heading-direction network of monarch butterflies could gain coding flexibility, similar to what has been reported in *Drosophila* ^14,15^.

### Heading coding in quiescence

Our methodological approach allowed us for the first time in any insect to discriminate between the influence of different idiothetic inputs on the heading coding, i.e. vestibular-like, proprioceptive and motor efferences coupled with reafferent signals. In addition to visual input from ring neurons, E-PG cells receive idiothetic input in the form of angular movement from P-EN cells ^20,21^. Since only 5% of our recorded neurons conjunctively encoded flight turns and heading, it is highly unlikely that our results originate from recordings of P-EN cells or neurons downstream of E-PG cells that themselves encode angular movements ^42^. Thus, E-PG cells represent the most plausible candidate recorded neurons in which remapping occurs in our study.

When quiescent butterflies were confronted with an ambiguous 2-sun stimulus, the heading-direction network was not able to unambiguously encode a particular heading. This is because visual cues dominated the heading coding in this situation, as also observed in our cue conflict experiment. Visual inputs are transmitted via the R2/4d ring neurons into the E-PG network ^14,17^. As ring neurons’ receptive fields are biased towards the fronto-lateral visual field in monarch butterflies ^43^, and also in fruit flies ^17^, the dominance of visual input is expected to bias the heading-direction network to the frontal visual field, which is what we observed in quiescent monarch butterflies. However, comparable to what has been found in cockroaches ^30^, the butterfly’ heading is not purely based on visual information. It also relies on vestibular-like idiothetic cues, such as mechanosensory inputs. As previously shown in fruit flies ^18^, mechanosensory signals from the antennae may be integrated via another set of ring neurons into the butterfly’ heading network to enable the actual heading to be sustained even in darkness. However, our conflict experiments demonstrate that in quiescent butterflies vestibular-like inputs would be less relevant than visual information from the sun stimulus, as allowing butterflies to fly while mechanically blocking their steering was not sufficient to induce a change in the preferred firing directions. This indicates that proprioceptive information from the flight apparatus is not responsible for changes in the neurons’ preferred firing direction. Thus, even in a flying butterfly deprived from turns, the sun stimulus, and thus the ring neurons activity, dominate the coding of heading in the central complex. The coding of heading based on visual inputs – and in the absence of steering commands – will therefore manifest in a body-related sun-bearing coding in quiescence and during fixed flight.

### Flight steering-dependent transformation into a sun compass

When the tethered flying butterflies were free to steer, we observed substantial shifts in the neurons’ preferred firing directions extending the heading-direction network to a 360° representation of angular space. Notably, the steering-induced switch to a 360° sun compass leads to an unambiguous heading coding during flight even when animals oriented with an ambiguous 2-sun stimulus. This requires the heading-direction network to relate the angular position of multiple cues with respect to the animal’s orientation, which results in an allocentric compass coding. We showed that this switch to a compass coding depends on the integration of motor efference copies and reafferent signals into the central complex. While the ring neurons dominate the coding in quiescence, our cue conflict experiment suggests that the heading coding in E-PG cells is mainly controlled by idiothetic signals from P-EN neurons during flight. The dominance of the P-EN input during flight was unexpected since results in *Drosophila* demonstrated that the coding of heading is dominated by visual information ^13,44^, similar to what has been found in vertebrates, including fish ^9^, birds ^45^, and rats ^10^. The differences observed in modality weighting in monarch butterflies could represent adaptations to a migratory lifestyle. Anatomical differences between the heading-direction systems of *Drosophila* and migratory desert locusts have been proposed to increase the robustness of the compass during long-distance navigation ^46^. Consistent with this idea, our results show that heading representation remains robust to sudden changes in the solar azimuth. Under these conditions, the heading-direction network primarily relies on idiothetic cues. The difference in modality weighting we observed in butterflies compared to previously reported in *Drosophila* may also be of a methodological nature. In *Drosophila*, the heading-direction network has been studied in head-fixed preparations with the visual stimulus moved according to intended angular movements ^13–15,18^. In our experiments, the monarch butterflies were actually steering which results in more naturalistic idiothetic inputs, including motor efference copies, reafferent signals, and vestibular-like signals than in head-fixed preparations.

Taken together, the transition of the heading-direction network into a sun-compass during flight requires the integration of multimodal information into the central complex. In nature, additional inputs, such as the time-compensation of the sun position ^4,47,48^ and/or the earth’s magnetic field ^49,50^, may provide additional cues to form a robust compass system that monarch butterflies use to efficiently maintain proper course orientation during their long migratory journey.

## Online Methods

### Animals

Monarch butterflies (*Danaus plexippus*) were ordered as pupae from Costa Rica Entomology Supply (butterflyfarm.co.cr) and kept in an incubator (HPP 110 and HPP 749, Memmert GmbH + Co. KG, Schwabach, Germany) at 25°C, 80% relative humidity and 12:12 light/dark-cycle conditions. After eclosion, the adult butterflies were transferred into a different incubator (I-30VL, Percival Scientific, Perry, IA, USA) at 25°C and 12:12 light/dark condition. Adults had access to 15% sucrose diluted in water *ad libitum*. In total, we tested 89 butterflies.

### Setup for simultaneous behavioral and electrophysiological recordings

To record neural activity in the butterfly brain, signals from custom-built tetrodes (see Tetrodes and Implantation) were passed through an electrode interface board (EIB-18; Neuralynx Inc., Bozeman, MT, USA) and an adapter board (ADPT-DUAL-HS-DRS; Neuralynx Inc., Bozeman, MT, USA) to a Neuralynx recording system (DL 4SX 32ch System, Neuralynx Inc., Bozeman, MT, USA). The neural activity was monitored using the software Cheetah (Neuralynx Inc., Bozeman, MT, USA). During recording, the heading direction of a butterfly was observed in a flight arena similar to the ones described before ^4,51^. The arena had an inner diameter of 32 cm and a height of 12 cm (Fig. 1A). A tungsten rod was attached to the butterfly’s thorax using a magnet (diameter = 3 mm; magnetic force = 4 N; Supermagnete, Webcraft GmbH, Gottmadingen, Germany) as well as dental wax (Article: 54895 Omnident, Rodgau Nieder-Roden, Germany) and was connected dorsally to an optical encoder (E4T miniature Optical Kit Encoder, US Digital, Vancouver, WA, USA) at the center of the arena. To avoid wrapping the tetrode wires around the tungsten rod, the animals’ rotatory movements were restricted to 358°. The heading direction was measured at a sampling rate of 100 Hz and with an angular resolution of 3°. These measurements were then digitized (USB4 Encoder Data Acquisition USB Device, US Digital, Vancouver, WA, USA). The animal’s heading was observed in the US Digital software (USB1, USB4: US Digital, Vancouver, WA, USA). A trigger signal at the onset of the behavioral recording was sent via an ATLAS analog isolator (Neuralynx Inc., Bozeman, MT, USA) and the adapter board to the Neuralynx recording system. This allowed us to align the behavioral data from the encoder with our neural recordings in the software Spike2 (version 9.0 Cambridge Electronic Devices, Cambridge, UK) after an experiment. The upper inner circumference of the flight arena was equipped with 144 RGB-LEDs (Adafruit NeoPixel, Adafruit Industries, New York, New York, USA) at an elevation of ~ 30° relative to the butterfly. One of these LEDs, when lit, provided a single green light spot as a stimulus (1.74 × 10^13^ photons/cm^2^/s and 1.2° angular subtense, as measured at the center of the arena). Insects interpret such a stimulus as the sun ^52–54^. The angular position of the green light spot was controlled with an Arduino MEGA 2560. An analog output from the Arduino was sent via the ATLAS analog isolator to the Neuralynx recording system and allowed us to align the stimulus presentation with the neural activity.

### Tetrodes and Implantation

To record neural activity in the butterfly brain, we adopted and modified the recording method that has been successfully established in honeybees to record learning related plasticity ^55–57^. A tetrode was built by waxing five (four recording wires and one differential wire) 18 cm long and 12.5 μm thin (10μm + insulation) copper wires (P155, Elektrisola, Reichshof-Eckenhagen, Germany) together. The tetrode was carefully threaded through two Pebax® tubes (each 2-4 cm in length; 0.026’ inner diameter; Zeus Inc, Orangeburg, SC, USA) that served as attaching points to mount the tetrode to a glass capillary. Tetrode and glass capillary were then attached to an electrode holder (M3301EH; WPI, Sarasota, FL, USA). The tetrode wires were connected to different channels of the EIB-18. A separate wire was connected to the grounding pin of the EIB-18 and served as grounding wire. To reduce the resistance of each electrode to about 0.1-1 MΩ, the tips of the tetrode wires were plated with gold (Elektrolyt Gold solution, Conrad Electronic SE, Hirschau, Germany). The resistance of each tetrode wire was measured using a nanoZ (Multi Channel Systems MCS GmbH, Reutlingen, Germany). To position the tetrode in the butterfly brain, the electrode holder (together with the EIB-18 and the tetrode) was connected to a micromanipulator (Sensapex, Oulu, Finland). For tetrode implantation, a monarch butterfly was horizontally restrained on a magnetic holder. The head was waxed to the thorax to avoid head movements during recording and flight. The head capsule was opened dorsally and fat and trachea on top of the brain were removed. To gain access to the central complex, the neural sheath on the dorsal brain surface was carefully removed using fine tweezers. The grounding wire was placed posteriorly in the head capsule. To identify the tetrode position, the tip of the tetrode was immersed in ALEXA 647 Hydrazide (A20502, Thermo Fisher Scientific GmbH, Dreieich, Germany) or ALEXA 568 Hydrazide (A10437, Thermo Fisher Scientific GmbH, Dreieich, Germany) diluted in 0.5 M KCl. Tetrodes were then inserted into the central brain while measuring neural signals with a sampling frequency of 30 kHz and a band-pass filter (600-6000Hz). One of the five tetrode wires was selected as a reference in the recording software and its signal was subtracted from the signals of the remaining four tetrode wires to obtain a differential signal. After finding neurons encoding the angular position of the sun stimulus, the tetrode and the grounding wire were held in place by adding a two-component silicone elastomer (Kwik-Sil, WPI, Sarasota, FL, USA). After the Kwik-Sil hardened (~ 1 hour), the butterfly was carefully unrestrained and connected to the end of the tungsten rod that was connected to the optical encoder. The tetrodes were carefully removed from the glass capillary and attached to a Pebax® tube to the tungsten rod. This setup allowed us to obtain stable neural recordings for several hours when the butterfly was either tethered flying or quiescent on a platform.

### Visualization of electrode tracks

After neural recording, the brain was dissected out of the head and fixated overnight in 4% formaldehyde at 4°C. After fixation, the brain was transferred into sodium-phosphate buffer (NaPi) and rinsed for 2 × 20 and 3 × 20 minutes in 0.1 M phosphate buffered saline (PBS) and PBS with 0.3% Triton-X (PBT), respectively. The brain was dehydrated with an ascending ethanol series (30% - 100%, 15 minutes each) and immersed with a 1:1 ethanol-methyl-salicylate solution for 15 minutes, followed by a clearing step in methyl-salicylate for at least 1 hour. The brain was embedded in Permount (Fisher Scientific GmbH, Schwerte, Germany) between two cover slips and scanned with a confocal microscope (Leica TCS SP2, Wetzlar, Germany) using a 10 × water immersion objective (HCX PL-Apo 10x/0.4 CS, Leica, Wetzlar, Germany). To visualize the position of the tetrode, we reconstructed the tetrode tracks together with subneurons of the central complex in 3D using the software Amira 5.3.3 (ThermoFisher, Germany). To visualize the tetrode positions from different experiments, we afterwards registered the tetrode position into the monarch butterfly standard central complex ^7^. We used an affine (12-degrees of freedom), followed by an elastic registration to transfer the neuropils of the individual central complexes into the corresponding neuropils of the standard central complex. The registration and deformation parameters were then applied to the tetrode reconstruction to visualize the tetrodes in one frame of reference.

### Spike sorting and spike shape analysis

Recordings were spike sorted with the tetrode configuration implemented in Spike2 (version 9.00, Cambridge Electronic Devices, Cambridge, UK). We used four spike detection thresholds (two upper and two lower thresholds). The highest and lowest thresholds were set to avoid large voltage deflections deriving from movement artifacts being misinterpreted as spikes. The time window for template detection was set to 1.6 ms. After spike-sorting, a principal component analysis (PCA) was used to evaluate and to redefine spike clusters, if necessary. Spike2 channels were exported as down-sampled Matlab files (3 kHz) and the remaining analysis was done with custom written scripts in MATLAB (Version R2019b, MathWorks, Natick, MA, USA). To analyze the spike shapes and to perform correlation analysis, the WaveMark channels containing the spike-waveforms were additionally exported as non-down-sampled Matlab files (30 kHz). To ensure that we recorded from the same neuron throughout the experiment, we compared the neuron-specific spike shapes across different recording periods. For each behavioral state e.g., *quiescent pre-flight, fixed flight, steering flight, quiescent post-flight*, the average spike shape was constructed by calculating and merging the mean spike shape from each recording electrode along the time axis. This resulted in a total time window of 6.4 ms (4 × 1.6 ms) or 232 points corresponding to a sampling rate of 30 kHz for each behavioral state. The average spike shape during pre-flight was compared to the average spike shapes measured during flight (e.g., *fixed flight*, *steering flight*), and after flight (*quiescent post-flight*). This resulted in at least two correlation coefficients per neuron. We used the lowest correlation coefficient of each neuron to compare it to a modelled distribution of correlation coefficients generated by the correlation between the neuron’s spike shape during pre-flight and all spike shapes obtained within one set of experiments (150 neurons for the state-dependent change experiments (Fig. 1,2,3,S2,S3); 55 neurons for pharmacological experiments (Fig. 5); 47 neurons for the passive rotation experiments (Fig. 6,S4); 37 neurons for the experiment using two suns (Fig 4)). The correlation with a randomly selected spike shape was repeated in total 1,000,000 times to obtain the distribution of correlation coefficients for each neuron. Only neurons that exhibited a statistically higher neuron-specific correlation coefficient than the modelled correlation coefficients (p < 0.05) were used for further analysis. 289 neurons passed this criterion and had an average correlation coefficient of 0.94 ± 0.08 (mean ± SD).

### Experiment testing state-dependent changes in the preferred firing direction

To examine the neural response to a green light spot during quiescence, a green light spot was moved 360° around the animal during recording while the butterfly was sitting on a platform (Compact Lab Jack, Inc, Newton, New Jersey, USA; *quiescent pre-flight*). The angular velocity of the sun stimulus was set to 60°/s which is commonly used to characterize compass neurons in restrained insects ^6,43,52^. The light spot was moved in clockwise and counterclockwise directions. While recording from the same neuron, the stimulus condition was repeated (360° movement of the sun stimulus around the butterfly) but this time while the animal performed a tethered flight in one locked direction (0°, the same heading direction as in quiescence, *fixed flight*). Afterwards, the position of the sun stimulus remained stationary at 0° while the tethered flying butterfly was free to rotate around its yaw axis (*steering flight*). In some experiments (n = 63 neurons), the butterfly was allowed to sit on the platform again to measure the neural response after flight (*quiescent post-flight*). To generate circular plots (bin size = 10°) showing the spatial tuning of recorded neurons and to calculate the preferred firing directions and vector strengths of each neuron, the CircStat ^58^ and CircHist ^59^ toolboxes of MATLAB were used. The neural sensitivity for coding the angular position of the sun stimulus was examined by testing if the neural response showed a non-uniform distribution (p < 0.05, Rayleigh test). If this was the case in one of the conditions, the preferred firing direction and the vector strength of a neuron was calculated.

### Pharmacological experiments

For the pharmacological experiments, we did not fix the tetrode’s position with Kwik-Sil to ensure that we still had access to the brain during the entire experiment.

For the pharmacological experiments, 10-20 μL of 500 μM chlordimeform (an octopamine agonist) diluted in saline was topically applied onto the brain while performing neural recordings in quiescent animals. We decided for a relatively high concentration of chlordimeform because the body dimensions of the monarch butterfly are larger than that of flies ^26,28^ and we had to ensure that enough chlordimeform diffused to the central complex (depth: 200-300μm). The neural sensitivity for coding the angular position of the light spot was measured before treatment and in 10-minute intervals from the onset of chlordimeform treatment. The stimulus was moved around the animal in the same way as described above for quiescence. The neurons were recorded for about 40-140 minutes (mean = 70 ± 29 minutes). The neural sensitivity to the stimulus was measured before treatment and compared with the last measurement.

### The influence of stimulus history and velocity on neural tuning

To test for the influence of stimulus history on neural tuning, the simulated sun was moved with a constant velocity of 60°/s in an unpredictable manner around a quiescent butterfly (*unpredictable stimulus history*). To test for changes in preferred firing directions associated with stimulus velocity, we also varied the velocity of the sun stimulus between 20-80°/s while it was moved in an unpredictable manner around the animal (*unpredictable stimulus velocity*; stimulus details are plotted in Fig. S3A). The preferred firing directions obtained in these experiments were compared to the neural response of a predictable moving sun stimulus (*quiescent*). At the end of each experiment, the preferred firing directions were also measured during flight (*steering flight*). The preferred firing directions in *quiescence* and during *steering flight* were compared with the preferred firing directions obtained in response to the unpredictably moving sun stimulus conditions (n = 42 neurons). A V-test (expected mean: 0°) was used to test if the change in preferred firing direction was similar between two conditions. If the V-test was significant (p < 0.05) this means that data were clustered around 0° implying that the preferred firing direction did not change between the tested conditions.

### Testing for coding of turning behavior

To test whether the recorded central-complex neurons encoded flight turns, we selected the flight sections in which the animal’s heading changed more than 9° from its actual heading direction. We set 9° as threshold for turns because the angular sensitivity of the optical encoder was 3° and deviations of ± 3° could represent variations in flight direction rather representing flight turns. We then examined the firing rate prior to and after the moment of a turn during a one-second window (500 ms prior and 500 ms after a flight turn). The neural activity during the turn was normalized to the firing rate 500 ms prior to the turn and was compared to the firing rate of sections in which the animal flew straight. Neurons were categorized as coding for flight turns if (i) changes in the normalized neural activity during flight turns were higher/lower than the changes occurring during periods of no turns (Wilcoxon p-test < 0.05) or (ii) if changes in the normalized neural activity during time windows of flight turns fitted a Gaussian distribution (> 0.6). 12 of the 150 (8%) neurons passed both criteria and were defined as neurons coding for flight turns. Only 7 of these 12 neurons additionally encoded the angular position of the light spot during flight.

### Passive rotation experiments

To test for the influence of idiothetic cues on neural tuning, we mounted a platform on a rotation stage (DT-50, PI miCos, Eschbach, Germany) at the center of the LED arena. The butterfly was then fixed on the platform and the animal was rotated around its yaw axis. The motions of the rotation stage were controlled by a custom written script in MATLAB. The neural response was first measured during *quiescence* as described above. Then, the butterfly was rotated clockwise and counterclockwise at 60°/s (*quiescence rotation*). During *quiescence rotation*, the sun stimulus was presented at 0° (in front of the animal at the beginning of rotation). Afterwards, the butterfly was rotated in the same way, but in the absence of any visual cue (*darkness rotation*). At the end of the experiment, the butterfly was unrestrained, and the neural response was measured while the butterfly was allowed to fly (*steering flight*). The directedness of the neural response during *rotation* and *darkness rotation* was represented by calculating the vector strength of the neural response. The preferred firing direction of each neuron (n = 16) was calculated for each behavioral state (*rotation, darkness rotation*, *quiescence*, *steering flight*). Only neurons that encoded the angular position of the simulated sun in all conditions were considered. If the preferred firing directions were similar across conditions, then the changes in preferred firing directions would cluster around 0° (V-test). Alternatively, if the changes were not found to cluster around 0°, they were tested for a uniform distribution (Rayleigh test: p > 0.05).

To test how the neural tuning changes if the animal is engaged in flight but is passively turned (instead of performing controlled steering), we performed similar experiments with the rotation stage mounted above the tethered animal. The tethered animal was then connected to the rotation stage and was passively rotated around its yaw axis (*rotation fixed flight*) while experiencing the simulated sun at 0°. The neural response of the neurons was compared to the responses recorded during quiescent situation (*quiescent pre-flight*) and when the animal was allowed to fly in one heading direction (*fixed flight*) while the sun stimulus was moved around the animal (at a velocity of 60°/s). After flight, the neural response was remeasured during quiescence (*quiescent post-flight*). The preferred firing directions of 17 neurons that encoded the angular position of the sun stimulus in all behavioral states were calculated and compared with the preferred firing direction during *quiescence* (pre-flight).

### Conflict experiments

To test the relevance of the sun-stimulus in different behavioral states, we performed the same experiments as described above (*rotation*, *steering flight*), with the exception that the position of the sun stimulus was switched to the opposite side of the arena (180°) after three clockwise and counterclockwise rotations (*rotation*) or after 2 minutes of flight (*steering flight*). Afterwards, the experiment was repeated with the sun stimulus located at 180° relative to its original position. The change of the preferred firing direction was compared between the two conditions. If the preferred firing direction changed by 180° (V-test, expected direction:180°), then the preferred firing directions were dominated by the change in the simulated sun’s position.

### Experiments with the two-sun stimulus

For the experiments with the two suns, two equally bright (1.74 × 10^13^ photons/cm^2^/s) green light spots were positioned 180° apart from each other (at 0° and 180° with respect to the initial orientation of the animal). The neural response was measured during *quiescence* (pre- and post-flight) while the two suns were moved clockwise and counterclockwise around the animal (angular velocity: 60°/s). Additionally, the neural response was measured while the tethered flying butterfly was able to steer in the presence of the two sun stimuli (*steering flight*). A weighted fit model was used to test if the neural response was described by a Gaussian (weight < 0.5; unimodal) or a sine-square function (weight > 0.5; bimodal) ^52^. 33 central-complex neurons were sensitive in each of the three tested conditions (*quiescent pre-flight, steering flight, quiescent post-flight*). The weights were statistically compared across the conditions according to a RM one-way ANOVA + Holm Sidák’s multiple comparison test (GraphPad Prism 9, GraphPad Software, San Diego, CA, USA).

### Data and codes

Matlab files with the calculated response parameters of the neurons together with the Matlab-scripts used for the analysis and Arduino scripts used for stimulus presentation are accessible from the Figshare Repository: https://….

## Acknowledgements

We thank Marie Dacke, Keram Pfeiffer, and Jan Ache for their helpful comments on our manuscript and Jan Ache for providing us with chlordimeform for pharmacological experiments. We thank Vivek Jayaraman, Manu Madhav and Flavio Rocces for discussing our results. We thank Keram Pfeiffer and Konrad Öchsner for their help in setting up the LED stimulus and Fabian Schmalz for introducing Neuralynx and spike sorting in *spike2*. In addition, we thank Sergio Siles (butterflyfarm.co.cr) and Marie Gerlinde Blaese for providing us with monarch butterfly pupae.

## References

1 Mouritsen, H. Long-distance navigation and magnetoreception in migratory animals. Nature 558, 50–59, doi:10.1038/s41586-018-0176-1 (2018).

2 Reppert, S. M., Gegear, R. J. & Merlin, C. Navigational mechanisms of migrating monarch butterflies. Trends Neurosci 33, 399–406, doi:10.1016/j.tins.2010.04.004 (2010).

3 Reppert, S. M., Guerra, P. A. & Merlin, C. Neurobiology of Monarch Butterfly Migration. Annu Rev Entomol 61, 25–42, doi:10.1146/annurev-ento-010814-020855 (2016).

4 Mouritsen, H. & Frost, B. J. Virtual migration in tethered flying monarch butterflies reveals their orientation mechanisms. Proc Natl Acad Sci U S A 99, 10162–10166, doi:10.1073/pnas.152137299 (2002).

5 Perez, S. M., Taylor, O. R. & Jander, R. A Sun compass in monarch butterflies. Nature 387, 29–29, doi:DOI 10.1038/387029a0 (1997).

6 Heinze, S. & Reppert, S. M. Sun Compass Integration of Skylight Cues in Migratory Monarch Butterflies. Neuron 69, 345–358, doi:10.1016/j.neuron.2010.12.025 (2011).

7 Heinze, S., Florman, J., Asokaraj, S., El Jundi, B. & Reppert, S. M. Anatomical basis of sun compass navigation II: the neuronal composition of the central complex of the monarch butterfly. J Comp Neurol 521, 267–298, doi:10.1002/cne.23214 (2013).

8 Finkelstein, A. et al. Three-dimensional head-direction coding in the bat brain. Nature 517, 159–164, doi:10.1038/nature14031 (2015).

9 Vinepinsky, E. et al. Representation of edges, head direction, and swimming kinematics in the brain of freely-navigating fish. Sci Rep 10, 14762, doi:10.1038/s41598-020-71217-1 (2020).

10 Taube, J. S. The head direction signal: origins and sensory-motor integration. Annu Rev Neurosci 30, 181–207, doi:10.1146/annurev.neuro.29.051605.112854 (2007).

11 Taube, J. S., Muller, R. U. & Ranck, J. B., Jr. Head-direction cells recorded from the postsubiculum in freely moving rats. I. Description and quantitative analysis. J Neurosci 10, 420–435 (1990).

12 Cullen, K. E. & Taube, J. S. Our sense of direction: progress, controversies and challenges. Nat Neurosci 20, 1465–1473, doi:10.1038/nn.4658 (2017).

13 Seelig, J. D. & Jayaraman, V. Neural dynamics for landmark orientation and angular path integration. Nature 521, 186–+, doi:10.1038/nature14446 (2015).

14 Fisher, Y. E., Lu, J., D’Alessandro, I. & Wilson, R. I. Sensorimotor experience remaps visual input to a heading-direction network. Nature 576, 121–125, doi:10.1038/s41586-019-1772-4 (2019).

15 Kim, S. S., Hermundstad, A. M., Romani, S., Abbott, L. F. & Jayaraman, V. Generation of stable heading representations in diverse visual scenes. Nature 576, 126–131, doi:10.1038/s41586-019-1767-1 (2019).

16 Hulse, B. K. & Jayaraman, V. Mechanisms Underlying the Neural Computation of Head Direction. Annu Rev Neurosci 43, 31–54, doi:10.1146/annurev-neuro-072116-031516 (2020).

17 Seelig, J. D. & Jayaraman, V. Feature detection and orientation tuning in the Drosophila central complex. Nature 503, 262–266, doi:10.1038/nature12601 (2013).

18 Okubo, T. S., Patella, P., D’Alessandro, I. & Wilson, R. I. A Neural Network for Wind-Guided Compass Navigation. Neuron 107, 9241–940 e918, doi:10.1016/j.neuron.2020.06.022 (2020).

19 Hardcastle, B. et al. A visual pathway for skylight polarization processing in Drosophila. Elife 10, ARTN e63225 doi:10.7554/eLife.63225 (2021).

20 Turner-Evans, D. et al. Angular velocity integration in a fly heading circuit. Elife 6, doi:10.7554/eLife.23496 (2017).

21 Green, J. et al. A neural circuit architecture for angular integration in Drosophila. Nature 546, 101–+, doi:10.1038/nature22343 (2017).

22 Weir, P. T. & Dickinson, M. H. Functional divisions for visual processing in the central brain of flying Drosophila. Proc Natl Acad Sci U S A 112, E5523–5532, doi:10.1073/pnas.1514415112 (2015).

23 Rosner, R., Pegel, U. & Homberg, U. Responses of compass neurons in the locust brain to visual motion and leg motor activity. J Exp Biol 222, doi:10.1242/jeb.196261 (2019).

24 Martin, J. P., Guo, P. Y., Mu, L. Y., Harley, C. M. & Ritzmann, R. E. Central-Complex Control of Movement in the Freely Walking Cockroach. Current Biology 25, 2795–2803, doi:10.1016/j.cub.2015.09.044 (2015).

25 Orchard, I., Ramirez, J. M. & Lange, A. B. A Multifunctional Role for Octopamine in Locust Flight. Annual Review of Entomology 38, 227–249, doi:DOI 10.1146/annurev.en.38.010193.001303 (1993).

26 Longden, K. D. & Krapp, H. G. State-Dependent Performance of Optic-Flow Processing Interneurons. J Neurophysiol 102, 3606–3618, doi:10.1152/jn.00395.2009 (2009).

27 Ache, J. M., Namiki, S., Lee, A., Branson, K. & Card, G. M. State-dependent decoupling of sensory and motor circuits underlies behavioral flexibility in Drosophila. Nature Neuroscience 22, 1132–+, doi:10.1038/s41593-019-0413-4 (2019).

28 Jung, S. N., Borst, A. & Haag, J. Flight activity alters velocity tuning of fly motion-sensitive neurons. J Neurosci 31, 9231–9237, doi:10.1523/JNEUROSCI.1138-11.2011 (2011).

29 Kathman, N. D. & Fox, J. L. Representation of Haltere Oscillations and Integration with Visual Inputs in the Fly Central Complex. Journal of Neuroscience 39, 4100–4112, doi:10.1523/Jneurosci.1707-18.2019 (2019).

30 Varga, A. G. & Ritzmann, R. E. Cellular Basis of Head Direction and Contextual Cues in the Insect Brain. Curr Biol 26, 1816–1828, doi:10.1016/j.cub.2016.05.037 (2016).

31 Shinder, M. E. & Taube, J. S. Self-motion improves head direction cell tuning. J Neurophysiol 111, 2479–2492, doi:10.1152/jn.00512.2013 (2014).

32 Coletta, S., Frey, M., Nasr, K., Preston-Ferrer, P. & Burgalossi, A. Testing the Efficacy of Single-Cell Stimulation in Biasing Presubicular Head Direction Activity. Journal of Neuroscience 38, 3287–3302, doi:10.1523/Jneurosci.1814-17.2018 (2018).

33 von Holst, E. & Mittelstaedt, H. Das Reafferenzprinzip - (Wechselwirkungen Zwischen Zentralnervensystem Und Peripherie). Naturwissenschaften 37, 464–476, doi:Doi 10.1007/Bf00622503 (1950).

34 Heisenberg, M. & Wolf, R. On the Fine-Structure of Yaw Torque in Visual Flight Orientation of Drosophila-Melanogaster. J Comp Physiol 130, 113–130, doi:Doi 10.1007/Bf00611046 (1979).

35 Webb, B. Neural mechanisms for prediction: do insects have forward models? Trends Neurosci 27, 278–282, doi:10.1016/j.tins.2004.03.004 (2004).

36 Kim, A. J., Fitzgerald, J. K. & Maimon, G. Cellular evidence for efference copy in Drosophila visuomotor processing. Nature Neuroscience 18, 1247–+, doi:10.1038/nn.4083 (2015).

37 Stackman, R. W., Golob, E. J., Bassett, J. P. & Taube, J. S. Passive transport disrupts directional path integration by rat head direction cells. J Neurophysiol 90, 2862–2874, doi:10.1152/jn.00346.2003 (2003).

38 Bender, J. A., Pollack, A. J. & Ritzmann, R. E. Neural activity in the central complex of the insect brain is linked to locomotor changes. Curr Biol 20, 921–926, doi:10.1016/j.cub.2010.03.054 (2010).

39 Suver, M. P., Mamiya, A. & Dickinson, M. H. Octopamine Neurons Mediate Flight-Induced Modulation of Visual Processing in Drosophila. Current Biology 22, 2294–2302, doi:10.1016/j.cub.2012.10.034 (2012).

40 Maimon, G., Straw, A. D. & Dickinson, M. H. Active flight increases the gain of visual motion processing in Drosophila. Nature Neuroscience 13, 393–U329, doi:10.1038/nn.2492 (2010).

41 Chiappe, M. E., Seelig, J. D., Reiser, M. B. & Jayaraman, V. Walking Modulates Speed Sensitivity in Drosophila Motion Vision. Current Biology 20, 1470–1475, doi:10.1016/j.cub.2010.06.072 (2010).

42 Shiozaki, H. M., Ohta, K. & Kazama, H. A Multi-regional Network Encoding Heading and Steering Maneuvers in Drosophila. Neuron 106, 126–+ (2020).

43 Nguyen, T. A. T., Beetz, M. J., Merlin, C. & El Jundi, B. Sun compass neurons are tuned to migratory orientation in monarch butterflies. Proc Biol Sci 288, 20202988, doi:10.1098/rspb.2020.2988 (2021).

44 Kim, S. S., Rouault, H., Druckmann, S. & Jayaraman, V. Ring attractor dynamics in the Drosophila central brain. Science 356, 849–U111, doi:10.1126/science.aal4835 (2017).

45 Ben-Yishay, E. et al. Directional tuning in the hippocampal formation of birds. Curr Biol, doi:10.1016/j.cub.2021.04.029 (2021).

46 Pisokas, I., Heinze, S. & Webb, B. The head direction circuit of two insect species. Elife 9, ARTN e53985 doi:10.7554/eLife.53985 (2020).

47 Sauman, I. et al. Connecting the navigational clock to sun compass input in monarch butterfly brain. Neuron 46, 457–467, doi:10.1016/j.neuron.2005.03.014 (2005).

48 Merlin, C., Gegear, R. J. & Reppert, S. M. Antennal Circadian Clocks Coordinate Sun Compass Orientation in Migratory Monarch Butterflies. Science 325, 1700–1704, doi:10.1126/science.1176221 (2009).

49 Guerra, P. A., Gegear, R. J. & Reppert, S. M. A magnetic compass aids monarch butterfly migration. Nat Commun 5, 4164, doi:10.1038/ncomms5164 (2014).

50 Wan, G., Hayden, A. N., Iiams, S. E. & Merlin, C. Cryptochrome 1 mediates light-dependent inclination magnetosensing in monarch butterflies. Nat Commun 12, 771, doi:10.1038/s41467-021-21002-z (2021).

51 Franzke, M. et al. Spatial orientation based on multiple visual cues in non-migratory monarch butterflies. J Exp Biol 223, doi:10.1242/jeb.223800 (2020).

52 el Jundi, B. et al. Neural coding underlying the cue preference for celestial orientation. P Natl Acad Sci USA 112, 11395–11400, doi:10.1073/pnas.1501272112 (2015).

53 Edrich, W., Neumeyer, C. & Helversen, O. V. Anti-Sun Orientation of Bees with Regard to a Field of Ultraviolet-Light. J Comp Physiol 134, 151–157, doi:Doi 10.1007/Bf00610473 (1979).

54 Rossel, S. & Wehner, R. Celestial Orientation in Bees - the Use of Spectral Cues. J Comp Physiol 155, 605–613, doi:Doi 10.1007/Bf00610846 (1984).

55 Strube-Bloss, M. F., Nawrot, M. P. & Menzel, R. Mushroom body output neurons encode odor-reward associations. J Neurosci 31, 3129–3140, doi:10.1523/JNEUROSCI.2583-10.2011 (2011).

56 Brill, M. F., Reuter, M., Rossler, W. & Strube-Bloss, M. F. Simultaneous long-term recordings at two neuronal processing stages in behaving honeybees. J Vis Exp, doi:10.3791/51750 (2014).

57 Strube-Bloss, M. F., Nawrot, M. P. & Menzel, R. Neural correlates of side-specific odour memory in mushroom body output neurons. Proc Biol Sci 283, doi:10.1098/rspb.2016.1270 (2016).

58 Berens, P. CircStat: A MATLAB Toolbox for Circular Statistics. J Stat Softw 31, 1–21, doi:DOI 10.18637/jss.v031.i10 (2009).

59 Zittrell, F., Pfeiffer, K. & Homberg, U. Matched-filter coding of sky polarization results in an internal sun compass in the brain of the desert locust. Proc Natl Acad Sci U S A 117, 25810–25817, doi:10.1073/pnas.2005192117 (2020).

