## Supplementary Information for "Flight-induced compass representation in the monarch butterfly heading network"

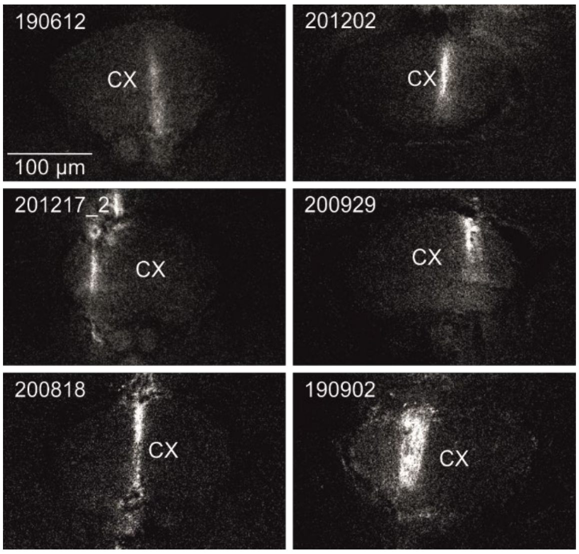


**Supplementary Figure 1. Visualization of tetrode tracks from six experiments.** Optical slices (z-slices) from the central body of the central complex showing the staining quality of tetrode tracks from some experiments. The tetrodes evidently run through the central body of the central complex (*CX*).

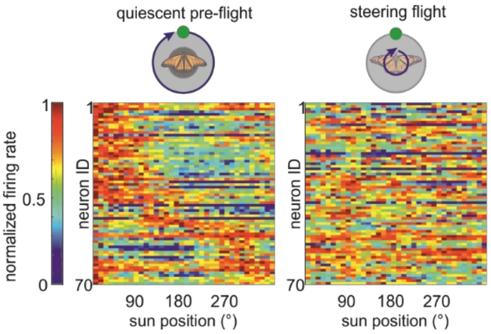


**Supplementary Figure 2. Differences in neural tuning between *quiescent pre-flight* and *steering flight*.** Neural tuning to the solar azimuth of all neurons showing angular sensitivity to the sun stimulus during *quiescent pre-flight* (*left*) and *steering flights* (*right*, n = 70 neurons). For reasons of comparison, neurons are ordered according to their preferred firing direction during *quiescent pre-flight*, along the y-axis.

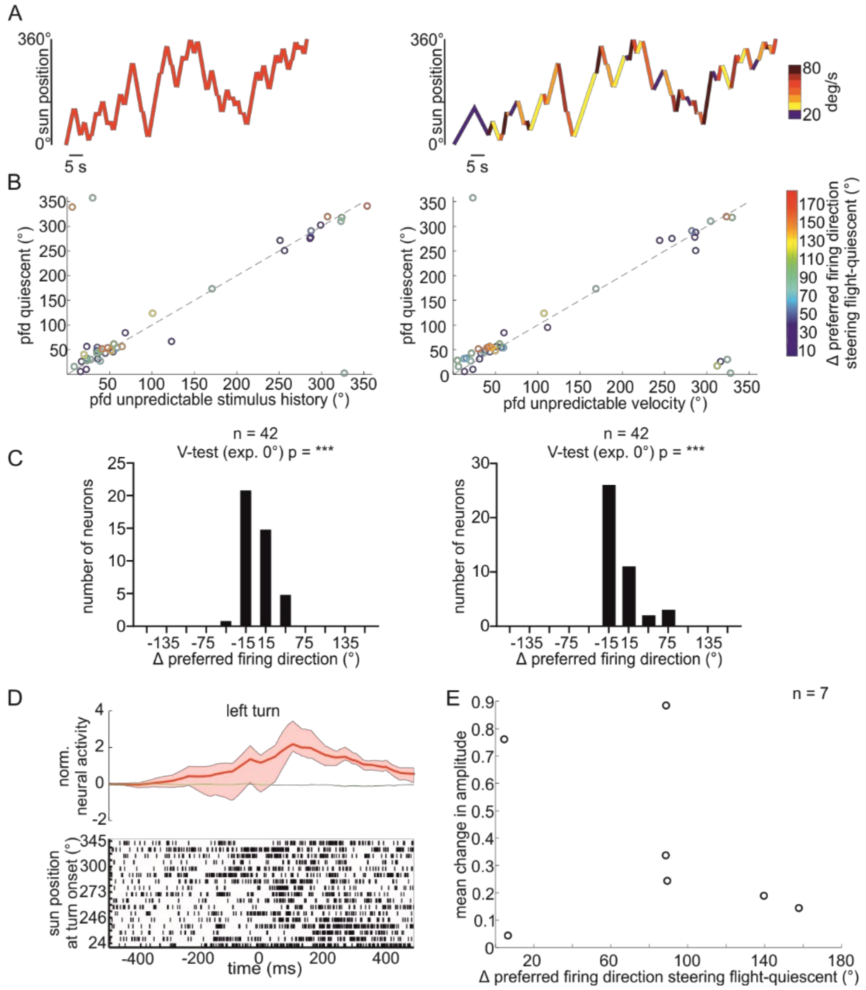


**Supplementary Figure 3. Stimulus history, stimulus velocity, and encoding turning behavior do not account for changes in preferred firing directions (pfd) observed during steering flights.** (**A**) (*Left*) Spatiotemporal dynamics of the stimulus that changes it's turning direction in an unpredictable manner (unpredictable stimulus history). (*Right*) Spatiotemporal dynamics of the stimulus that changes its turning direction *and* angular velocity (ranging from 20°-80°/s) in an unpredictable manner. (**B**) (*Left*) Scatter plot showing the relationship between the preferred firing direction during quiescence in which the stimulus was rotated with a constant velocity of 60°/s in full turns around the butterfly and the preferred firing direction calculated when the sun stimulus movement direction was unpredictable. Afterwards, the preferred firing direction was measured as the butterfly performed a steering flight. Color coded are the neural tuning shifts observed during steering flight. (*Right*) Same as in the left scatter plot but comparing the preferred firing directions between quiescent and unpredictable stimulus velocity. The preferred firing directions were similar between the compared stimulus conditions and neurons with large tuning shifts during steering flights (orange to red colored dots) do not show strong tuning shifts across variations in stimulus history or velocity. (**C**) Distribution of changes in preferred firing direction between quiescent and unpredictable stimulus history (*left*) and quiescent and unpredictable stimulus velocity (*right*). ***: $p < 0.001$; V-test (tested against 0°). (**D**) (*Top*) Sliding average shows the mean response (± interquartile range) to left turns for one example neuron. The averaged response at periods in which no turns occurred are plotted as green line. (*Bottom*) Raster plot of the example neuron in which the trials were ordered along the starting position of left turns. (**E**) Scatter plot of the response strength to flight turns, indicated by the mean change in response amplitude versus the shifts in preferred firing direction during steering flights. Only 7 out of the 150 recorded neurons responded to flight turns and also encoded the sun position during steering flight. No correlation between the neural tuning shifts during steering flights and the response strength to flight turns was observed which shows that the neural shifts cannot be explained by coding of turns.

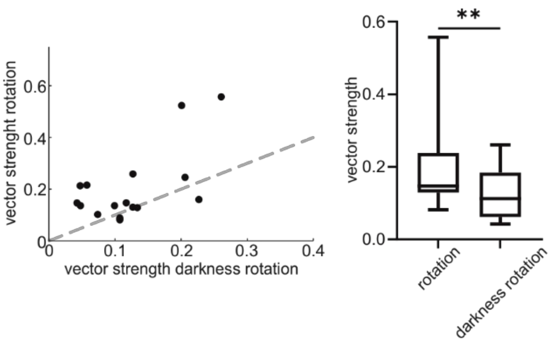


**Supplementary Figure 4. Spatial tuning of heading-direction cells is broader during passive rotations in darkness than in the presence of the sun stimulus.** Scatter plot (*left*) and boxplots (*right*) show the decrease in vector strength of the spatial tuning to the sun stimulus during passive rotations in darkness than during passive rotations in the presence of the sun stimulus. **: $p < 0.005$; Wilcoxon matched-pairs signed rank test.

**Supplementary Movie 1.** Demonstration of the stimulus when recording from a quiescent butterfly that was stimulated with a sun stimulus moving along a 360° circular path around the animal. For visualization the stimulus intensity was higher compared to when performing the experiments.

**Supplementary Movie 2.** Demonstration of the stimulus when recording from a *fixed flight* while the sun stimulus was moved along a 360° circular path around the animal. For visualization the stimulus intensity was higher compared to when performing the experiments.

**Supplementary Movie 3.** Demonstration of the stimulus when recording from a *steering flight* in which the flying butterfly was allowed to steer relative to a static sun stimulus. For visualization the stimulus intensity was higher compared to when performing the experiments.

**Supplementary Movie 4.** Demonstration of the stimulus when recording from a quiescent butterfly that was stimulated with a sun stimulus moving with a constant angular velocity (60°/sec) in unpredictable directions along a 360° circular path around the animal. For visualization the stimulus intensity was higher compared to when performing the experiments.

**Supplementary Movie 5.** Demonstration of the stimulus when recording from a quiescent butterfly that was stimulated with a sun stimulus moving with a varying angular velocity (20-80°/sec) in unpredictable directions along a 360° circular path around the animal. For visualization the stimulus intensity was higher compared to when performing the experiments.

**Supplementary Movie 6.** Demonstration of the stimulus when recording from a quiescent butterfly that was rotated in presence of a sun stimulus (*left*) and in darkness (*right*). For visualization the stimulus intensity was higher compared to when performing the experiments.

**Supplementary Movie 7.** Demonstration of the stimulus when recording from a fixed flying butterfly that was rotated by a dorsal rotation stage while a static sun stimulus was presented.

For visualization the stimulus intensity was higher compared to when performing the experiments.
